# Robust Harmonization of Microbiome Studies by Phylogenetic Scaffolding with MaLiAmPi

**DOI:** 10.1101/2022.07.26.501561

**Authors:** Samuel S Minot, Bailey Garb, Alennie Roldan, Alice Tang, Tomiko Oskotsky, Christopher Rosenthal, Noah G Hoffman, Marina Sirota, Jonathan L Golob

## Abstract

Microbiome science is difficult to translate back to patients due to an inability to harmonize 16S rRNA gene-based microbiome data, as differences in the technique will result in *different* amplicon sequence variants (ASV) from the *same* microbe. Here we demonstrate that placement of ASV onto a common phylogenetic tree of full-length 16S rRNA alleles can harmonize microbiome studies. Using *in silico* data approximating 100 healthy human stool microbiomes we demonstrated that phylogenetic placement of ASV can recapitulate the true relationships between communities as compared closed-OTU based approaches (Spearman R 0.8 vs 0.2). Using real data from thousands of human gut and vaginal microbiota, we demonstrate phylogenetic placement, but not closed OTUs, were able to group communities by origin (stool vs vaginal) without being confounded by technique and integrate new data into existing ordination/clustering models for precision medicine. This enables meta-analysis of microbiome studies and the microbiome as a biomarker.

## Main

With the development of high-throughput sequencing, a myriad of studies have associated the human microbiome (the collection of microbes that live within and upon us) with health and disease ^1–6^. As of 2022, at least 2000 BioProjects in the NCBI sequence read archive (SRA) contain human microbiome data spanning over 150,000 individual specimens. Due to challenges with recruiting and retention, microbiome studies are often conducted at a single center and with limited numbers of participants. A complication has arisen as a result: studies of how the microbiome relates to the same scientific question frequently fail to reproduce consistent observed associations^7^. For example, multiple studies have associated the human gut microbiome with the efficacy of immune checkpoint inhibitor therapy, with each study finding a *different* set of bacterial species that associate with a response^8–12^. A similar challenge has arisen with the vaginal microbiome and preterm birth^13^. This has limited the translation of microbiome science to improved clinical care. The inconsistency of smaller single-center studies is not a unique problem for microbiome studies; similar challenges exist for studies associating with transcription, genetics, and epigenetics. With those ‘omics studies, meta-analysis by combining raw data at the sequence- or feature-level can overcome the challenges of small and single site studies^14^. But a fundamental technical challenge has blocked the combination of microbiome studies, particularly those that target the 16S rRNA gene ^6^.

The dominant technique in microbiome science has been amplicon sequencing of a hypervariable region of a taxonomically informative gene such as the 16S rRNA gene. There are nine hypervariable regions in the 16S rRNA gene, each of a size suitable for current high-throughput sequencing platforms. The 16S meta-analysis challenge arises when studies target different variable regions, or even the same variable regions but with differences in the PCR primers, PCR conditions, sequencing library preparation, and the sequencer itself. These technical differences result in the *same* underlying allele being detected as a *different* amplicon sequence variant (ASV), and thus not able to be directly combined and compared (**Figure 1A**). Thus, some normalization must occur to convert observations of individual sequences or inferred sequences (ASV) into a set of features that are comparable across studies by partitioning related ASVs into bins. Several approaches have emerged for binning reads, generally relying upon some outside reference. Common approaches include closed reference operational taxonomic units (cOTU) and projection to taxonomy (e.g., quantifying each family of microbes present). The resulting feature sets may then be used to compare sequence populations observed in two or more specimens using a range of methods, some non-hierarchical (e.g, Shannon entropy), others explicitly hierarchical (e.g., UniFrac).

**Figure 1:**
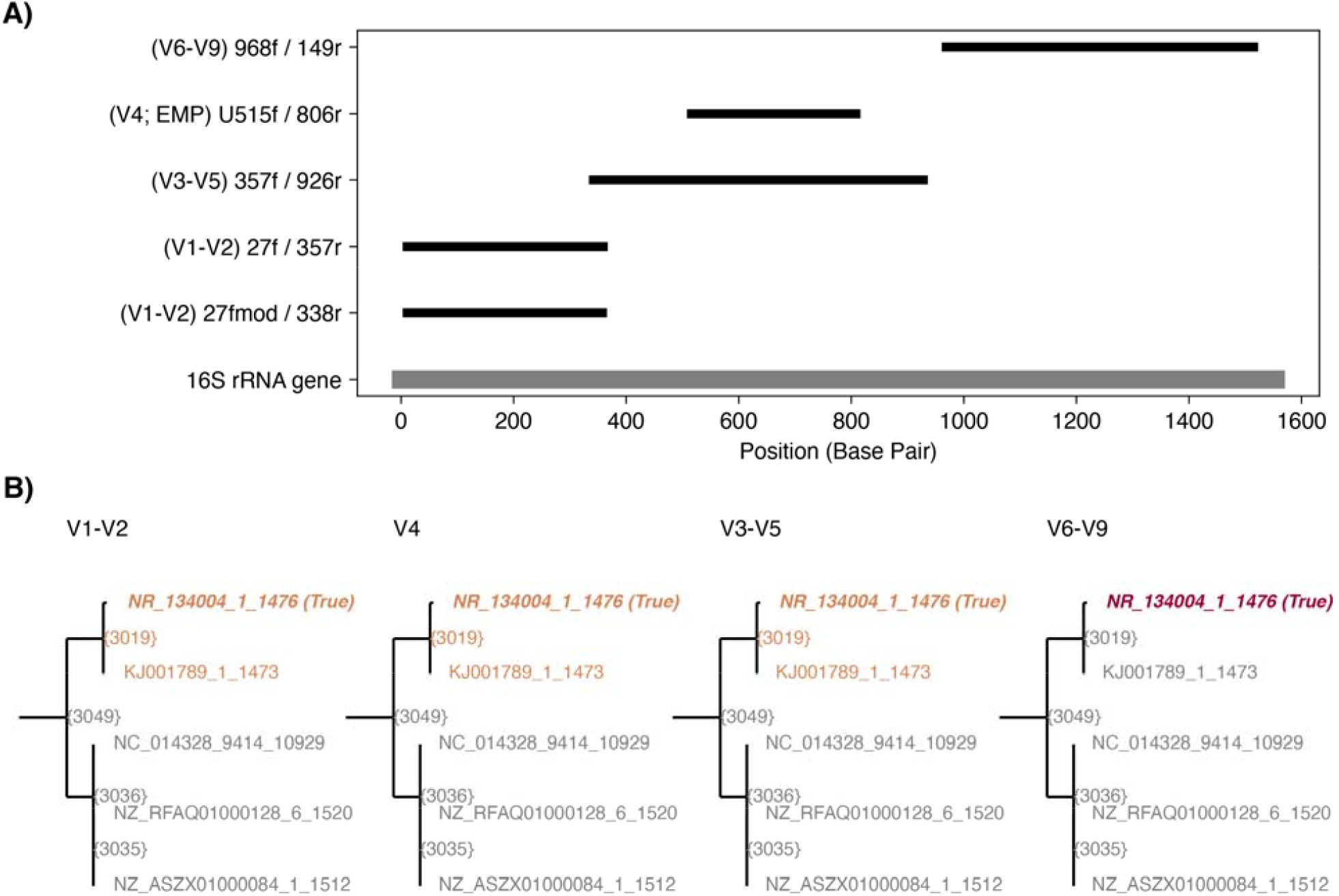
Non-overlapping sequences generated by 16S Variable Region Primers are placed on the same subclade of a full-length phylogenetic tree. (A) Primers targeting four different regions of the 16S rRNA gene: V1-V2, V4, V3-V5, and V6-V9 have largely non-overlapping positions within the full 16S rRNA gene. (B) A subclade of the reference tree is shown, with nodes of this subclade colored based upon the likelihood ratio of the given ASV placing at that node, with grey no likelihood. Despite each ASV having a *different* sequence they all phylogenetically place to the same small subclade of the phylogenetic tree which contains the true source allele, with the V6-V9 ASV containing sufficient entropy to entirely be placed on the true allele.

In cOTU generation each experimentally-derived amplicon sequence is aligned against a reference database of full-length 16S sequences^15^. Amplicon sequences that align best to the same reference sequence are grouped together. This technique is highly dependent on how well matched the reference is to the microbial communities being studied. Amplicon sequences without a good matching reference end up lost in this approach. Likewise, some amplicon sequences can have multiple nearly identically scored alignments to reference sequences, particularly when a very broad reference set is used. Adjudicating those nearly identically scoring alignments is a difficult challenge, and can lead to sequences from the same underlying true 16S rRNA allele being assigned ambiguously to multiple reference sequences.

Annotation of 16S rRNA gene variable region sequence variants with taxonomy, and then grouping read-counts at a selected (often Family or higher) taxonomic level is a common tactic (e.g. ^16,17^). Taxonomic assignments to variable region amplicon sequences are limited by the generally poor reliability of taxonomic assignments, particularly at more granular ranks (e.g. species of genus). As we have previously described, many classifiers can confidently misidentify the true underlying genus or species^18^. Further, divergence between phylogeny and taxonomy is a fundamental limit of this approach.

Phylogenetic placement methods for binning sequences or ASVs “place” sequences onto an existing phylogenetic tree ^19^, thereby mapping sequence observations onto tree-derived features such as specific edges of the tree graph. These methods have a number of advantages. Robust methods are available for accommodating and expressing uncertainty deriving from sequence variation^20^. An advantage over taxonomic methods and OTU-based methods limited to well-characterized taxa is that granular bins may be defined even for poorly defined taxonomic regions (e.g., environmental specimens, complex anaerobic environments such as the gut, etc.). Features are intrinsically hierarchical, so bins of varying granularity may be easily derived. The feature hierarchy is derived explicitly from relevant sequence data, in contrast to a taxonomy, which may either be discordant with sequence-based relationships, or define categories that are indistinguishable using available sequence data.

Summary metrics (e.g., Shannon alpha diversity) can address some of the current technical limitations. A robust example of this approach is the clinically relevant association of a loss of alpha diversity and risk for *C. difficile* infection. While somewhat buffered against the technical differences (variable region, primers, polymerase, sequencer), systemic biases can be introduced by things like differences in read depth and/or entropy in a given variable region. While relative differences in alpha diversity measures may be reproduced across studies, absolute values are not generally comparable across studies with different experimental parameters. Thus, we have also come to appreciate how much subtle technical biases affect estimates of alpha diversity ^21^.

Whole genome shotgun sequencing (WGS) is an alternative technique for microbiome studies, but with its own set of analytic challenges and opportunities^22,23^. The semi-random priming of reads eliminates some but not all of the cross study comparability problems between studies, as it does not eliminate differences in sequencers, library preparations, and sequencing depths. It also remains unclear how to integrate WGS and 16S rRNA data into one cohesive data set.

Here we demonstrate a technique that places 16S rRNA gene variable region amplicon sequence variants (ASVs) onto a common phylogenetic tree of *full length* 16S rRNA alleles and use tree-based methods for comparison among populations. We observe that this technique successfully corrects for the technical differences in 16S sequencing methods across studies, retains more true entropy, correctly combines different ASVs from the same underlying allele, and provides robust estimates of pairwise phylogenetic distance, alpha-diversity, and taxonomy. We have extended this approach to successfully integrate 16S gene and WGS-based microbiome studies, scaling up to tens of thousands of specimens on routinely available computational resources. This technique is available as a portable and reproducible containerized *Nextflow* workflow (MaLiAmPi), and immediately applicable to meta-analysis of 16S rRNA-gene based microbiome studies as well as clinical translation of extant studies.

## Results

### Working principles and the core approach of phylogenetic placement

The core approach implemented in MaLiAmPi is to generate a common reference phylogenetic tree that the amplicon sequence variants (ASV) are related to via phylogenetic placement. There are four steps: (1) generation of ASVs; (2) selection of a repository of full-length 16S rRNA alleles; (3) generation of a reference package including a phylogenetic tree of full-length 16S rRNA alleles from the repository that match the ASVs ; and (4) placement of the ASVs onto the reference package phylogenetic tree.

The core and motivating observation of this approach is that ASVs generated from the *same* underling 16S rRNA allele but with primers targeting *different* variable regions of the 16S rRNA gene will end up phylogenetically placed in the same small subclade of the reference phylogenetic tree, when the phylogenetic tree is well matched to the ASVs and comprised of full-length 16S rRNA alleles (**Figure 1B**). We then conjectured that clustering ASVs based on phylogenetic distance of their placements on a common reference phylogenetic tree would allow us to generate a count of reads per *<u>phylotype</u>* or groups of ASVs all of which were likely from the same (or very similar) underlying 16S rRNA allele.

### Successful combination of 16S rRNA amplicons in sequencing-error-free simulated data

We started with the idealized situation in which there is no sequencing error, and each amplicon’s sequence was available error-free and full-length to compare the performance of closed OTU generation versus phylotype-based grouping of amplicon sequence variants (ASV). The *in silico* amplicons (and full-length 16S rRNA alleles for comparison) were (i) dereplicated at 100% identity; processed with (ii) QIIME in a closed-reference approach against the GreenGenes 97% identity reference, with a goal of 80% similarity OTUs; or (iii) phylogenetically placed on common reference tree with MaLiAmPi, with ASVs grouped into phylotypes.

Ideally, a normalization technique would retain true community-to-community differences while eliminating false differences introduced by technical details (primers, PCR conditions, sequencer, etc.). In **Figure 2A** we use ordination plots to show the true relationship between the 100 simulated communities, note that with dereplication or closed-OTUs these relationships are lost, with the primer selected the dominant driver of clustering. Narrowing in on five randomly selected communities (**Figure 2B**), we can see that only with phylotype normalization does the representation of the same community with different primers tend to closely cluster into one group. As these are synthetic communities, we know the ‘true’ pairwise distance between them. Phylotype pairwise Bray Curtis distance was strongly correlated to the true pairwise distance between the simulated communities (Spearman R of 0.8) as compared to that estimated by closed OTUs (Spearman R 0.2) or dereplicated ASV sequences (Spearman R of 0) (**Figure 2C**).

**Figure 2:**
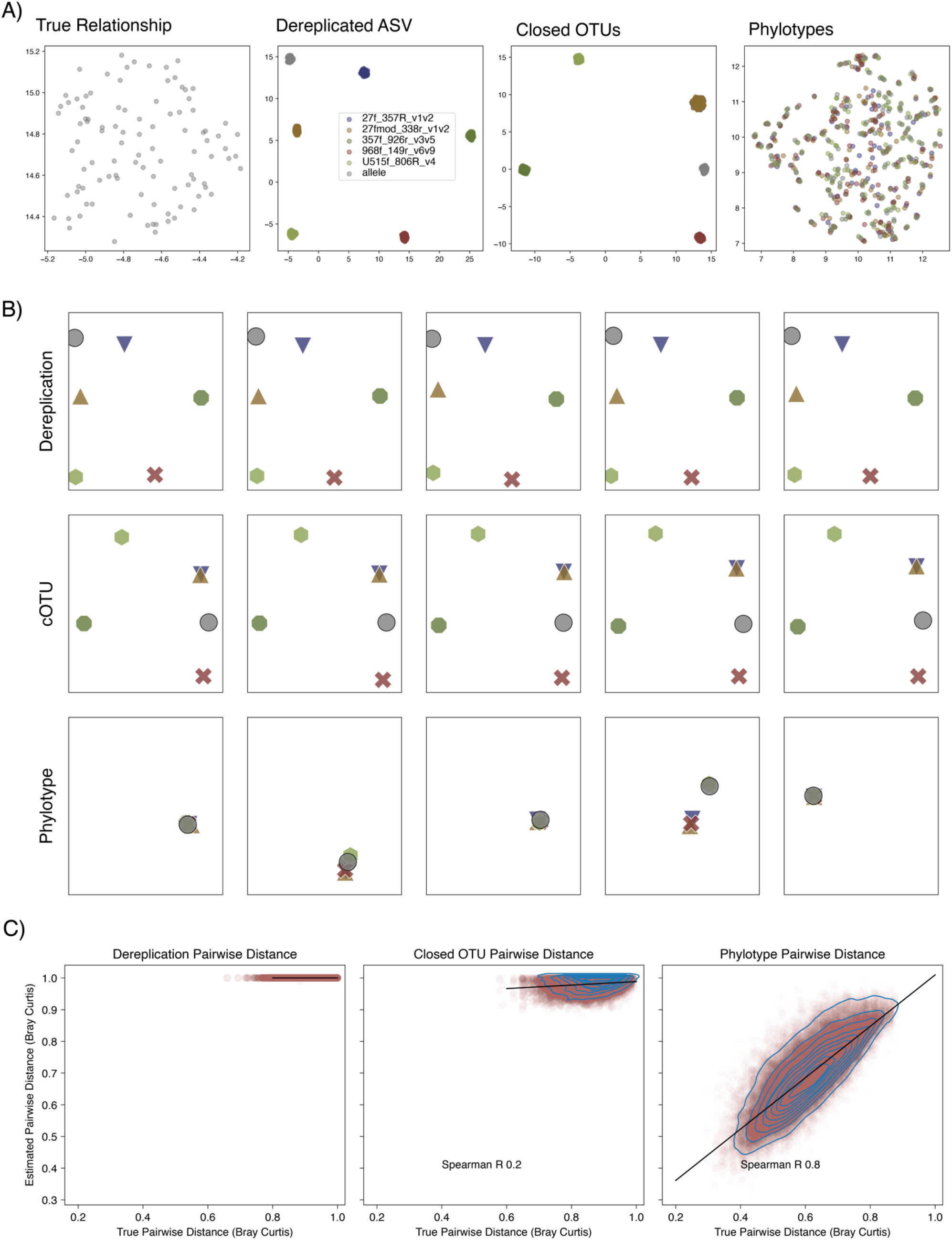
UMAP Ordination based on pairwise Bray-Curtis distance between 100 simulated microbial communities without sequencing error simulation. A) Each circle is a microbial community. The left most block is the true relationship between communities. Relationships based on Bray-Curtis distance of strictly dereplicated ASV (second to the left), closed OTUs (third from the left) and phylotypes (right), with grey representing full-length alleles and each color representing as amplified by a specific primer pair. Only phylotypes correctly estimate the true relationships between communities. B) UMAP ordination of five randomly selected communities based upon Bray Curtis distance of feature-counts, with each color and symbol representing a primer set used for amplification, where ideally these will all overlap for the a given community. Only phylotypes (and not closed OTUs or dereplicated ASVs) group by the source community rather than primer. C) A correlation between the ‘true’ Bray-Curtis distance between communities (x-axis) versus as estimated (y-axis) after dereplication, closed-OUT generation, or grouping into phylotypes. Ideally these would be perfectly correlated. R values reported are Spearman correlation coefficients.

### Successful correction of sequencer-introduced variation with DADA2 error deconvolution and phylogenetic placement

To consider the effect of sequencing error we simulated sequencing with two distinct platforms (454 pyrosequencing and Illumina MiSeq); for Illumina MiSeq, we applied three different empirically derived error models (to simulate the effect of per-batch and per-sequencer effects even when selecting the same primers and PCR conditions), the MsV1 model built into the ART package, a model derived from the University of Michigan Microbiome Core, and from PRJNA701859 sequenced at a microbiome center in Germany. While 454 pyrosequencing is no longer commercially available, a large amount of microbiome study data available in public repositories was generated on that platform. These simulated sequenced reads were processed with: (I) DADA2 as error-corrected amplicon sequence variants (ASV); (II) QIIME in a closed-reference approach against the GreenGenes 97% identity reference, with a goal of 80% similarity OTUs; (III) phylogenetically placed on common reference tree with MaLiAmPi, with ASVs grouped into phylotypes. The results (**Figure S1**) are extremely similar to what we observed with error-free reads: Exact sequence variants as deconvoluted from DADA2 and closed-reference OTU generation were able to overcome sequencer-introduced variation, but largely not the differences from different amplifying primers. In contrast, phylogenetic placement, and then derivation of phylotype-counts was able to properly group together by community rather than primer. This is reflected in the correlation between the true Bray Curtis pairwise distance (as estimated using the same primers and error models) versus ‘normalized’ Bray Curtis distance for the same pairs of communities but when using different primers and error models to generate the data, where the Spearman R was zero for dereplicated ASVs, 0.4 for closed OTUs, and 1 after phylogenetic normalization.

### MaLiAmPi phylogenetic placement integrates thousands of human microbiome specimens from multiple studies while retaining distinctions between vaginal and gut microbiota

We next obtained publicly available raw read data from studies of the vaginal microbiome during pregnancy and the healthy human gut (**Table 1**). We selected studies that would be particularly favorable to non-phylogenetic approaches: targeting similar variable regions of the 16S rRNA genes (when considering a specific body site) and with similar sequencing technology. Still, the reads-per-specimen and other technical aspects varied within this curated set of studies. This pilot involved over 5,000 specimens and over two million reads and was accomplished within modest computational resources (32GB of RAM; 12 core CPU).

**Table 1:** Sources of publicly available real-world data of human gut and vaginal microbiota.

| SRA Project | Body Site | Number of Specimens | Number of Reads | Variable Region | Sequencer | Study Location |
| --- | --- | --- | --- | --- | --- | --- |
| PRJEB14839 | Gut | 1,553 | 403,184 | V3-V5 | Illumina | Toronto Canada |
| PRJEB31801 | Gut | 8 | 10,346 | V3-V4 | Illumina | France |
| PRJNA418115 | Gut | 98 | 24,356 | V4 | Illumina | Michigan, USA |
| PRJNA607556 | Gut | 33 | 32,232 | V4 | Illumina | Minnesota, USA |
| PRJNA663047 | Gut | 50 | 6,952 | V3-V4 | Illumina | China |
| PRJNA701859 | Gut | 814 | 1,585,590 | V3-V4 | Illumina | Germany |
| PRJEB21325 | Vaginal | 157 | 10,887 | V1-V2 | Illumina | UK |
| PRJEB30642 | Vaginal | 253 | 5,059 | V1-V2 | Illumina | UK |
| PRJNA393472 | Vaginal | 2,175 | 173,624 | V1-V3 | Illumina | California, USA |
| **Total** |  | 5,141 | 2,252,230 |  |  |  |

### Disambiguating body site from experimental batch across published vaginal and gut microbiome datasets

Our first test was to see if phylotype normalization could result in specimens from different studies grouping by the site from which they were collected while minimizing the effect of the study techniques. Bray-Curtis pairwise distance was calculated from dereplicated ASV and phylotype counts. Ideally this distance would be strongly related to the site from which a specimen was collected, but not the study protocol under which it was collected. We used the ANOSIM statistic which takes a distance matrix as the independent data, a grouping as the dependent variable and results in a test statistic that is bounded from -1 (perfectly anticorrelated) to +1 (perfectly correlated) with zero being no correlation^24^. Ideally the ANOSIM statistic would be zero when grouping by project within each body site (assuming the study populations are biologically identical) and one when grouping by site of collection (assuming there is no true overlap between the vaginal and gut microbiome). Phylotype-counts come close to the ideal, with an ANOSIM R of 0.97 with the body site of collection, and only 0.17 or -0.07 to project for gut and vaginal microbiota respectively (**Figure 3**). This is reflected in the UMAP ordinations, where only the phylotype counts result in grouping by body site rather than project (**Figure 3 and S2**).

**Figure 3:**
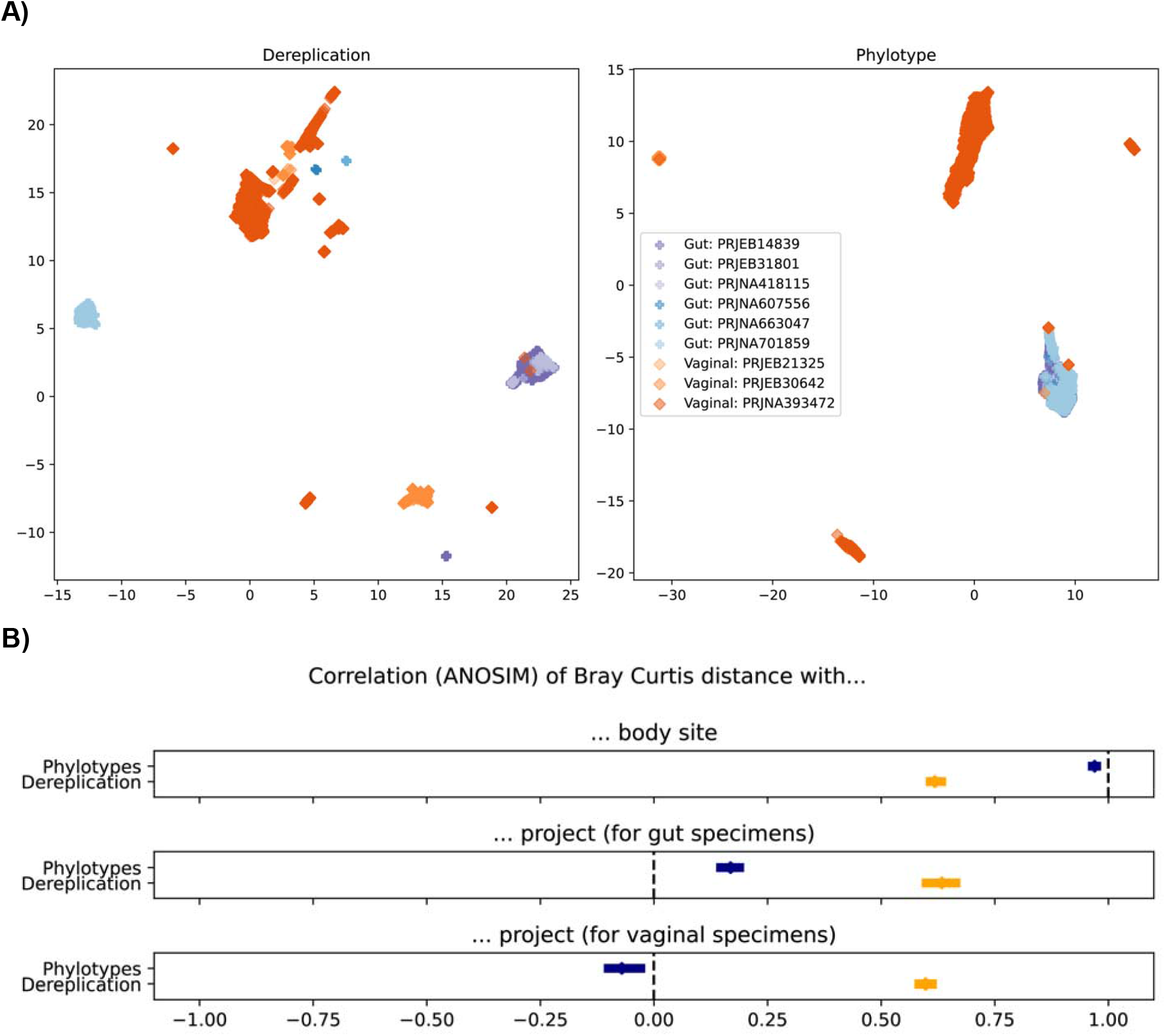
**A)** UMAP ordination based on Bray-Curtis pairwise distance between six gut (purples) and three vaginal (oranges) human microbial communities as estimated by dereplicated ASV counts (left) or phylotype counts (right). **B)** Correlation between the Bray-Curtis pairwise distance between microbiome specimens as estimated by phylotype (navy) or dereplication (orange) counts to the body site (top) from which the specimen was collected or the project that collected the gut (middle) or vaginal (bottom) microbiome specimen. Dashed vertical lines are the ideal outcome. Error bars are the 95% confidence interval by bootstrapping of the ANOSIM R statistic.

### Assigning human gut microbiome specimens to consistent ecotypes

It has been previously noted that the healthy human gut microbiome clusters into ecotypes that in turn can relate to health, such as the *Firmicutes* / *Bacteroides* ratio and obesity^25^. Generalizing these findings has been difficult^26^ in part due to the technical challenges with integrating microbiome data with established techniques and the reliability of taxonomic assignments. Using phylotype counts from the gut microbiome studies in **Table 1** we were able to consistently group specimens into two ecotypes; these ecotypes were in a similar proportion (1:4) across all six studies observing the healthy human gut microbiome (**Figure 4A**).

**Figure 4:**
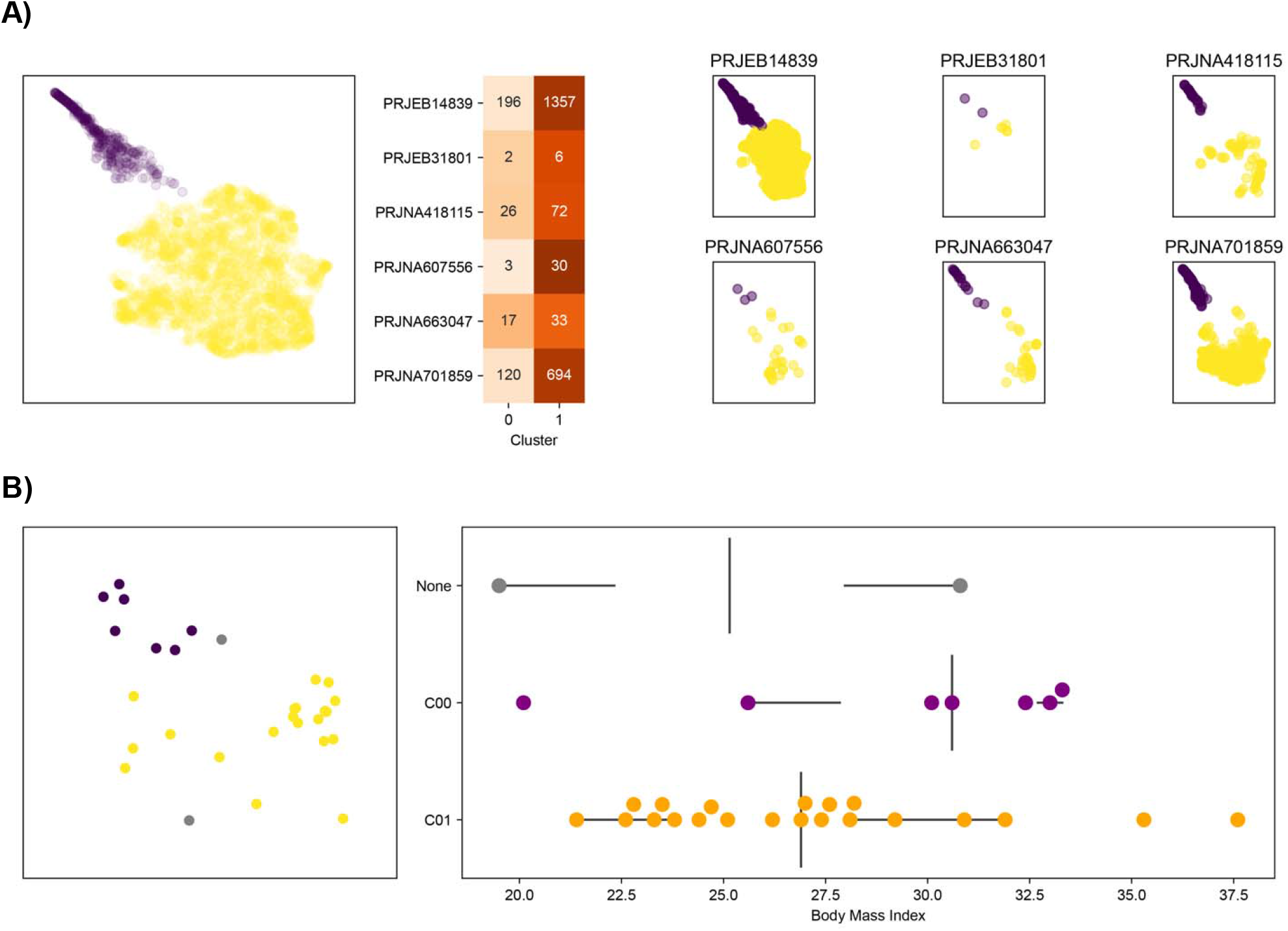
Assignment of gut microbiome specimens to two ecotypes that are consistent across studies. (A) Human gut microbiota specimens from six distinct studies were assigned to two clusters (purple or yellow) based on phylotype count Bray-Curtis distance after UMAP ordination and clustering by HDBSCAN. The proportion of healthy human gut specimens assigned to a cluster was consistent across studies. (B) We applied the ordination and clustering models to a different dataset relating the gut microbiome to body mass index. On the left, purple are specimens assigned to cluster 0, orange are assigned to cluster 1 and grey could not be assigned to a cluster. On the right are the body mass index values recorded stratified by the assigned cluster.

We then applied UMAP (ordination) and HDBSCAN (clustering) models generated from the phylotype counts from the six surveys of healthy adults to a *different* study of thirty adults that related the gut microbiome to body mass index (SRA BioProject PRJEB4203). Despite this study using 454 Pyrosequencing rather than the Illumina sequencing for the six surveys of the healthy human gut used to fit the models, we were able to integrate all thirty of the PRJEB4203 specimens into the existing phylotypes and ordination, and assign 28 of the thirty to an existing cluster. We noted that gut microbiota in cluster zero tended to have higher BMI (**Figure 4B**).

## Discussion

Revisiting prior ‘omics studies has proven a fruitful way to improve patient care. For example, it is now common practice to use genomics data to personalize and optimize cancer treatment regimens, significantly improving outcomes for patients^27,28^. Similarly, transcriptional^29^ and epigenomics studies are being combined and revisited with newer machine-learning techniques with an eye towards drug repurposing and personalized medicine. Facilitating these efforts are a very clear and intrinsically generalizable set of features, such as SNPs (genomics), loci (epigenomics), and genes (transcriptomics). Microbiome studies have lacked such a clear and generalizable underlying feature. Closed OTUs and taxons have been attempted when integrating microbiome studies, but both have fundamental limits that we have redemonstrated here or in previous studies^18^. Thus, a lack of a robust and generalizable feature has been a core limitation of microbiome science. It has left the field unclear of how to apply the findings of a study to other studies of the same clinical question and to an individual patient and use the microbiome as a biomarker (as is done with genomics data in cancer treatment).

Here we demonstrate the ability of phylogenetic placement of amplicon sequence variants from 16S rRNA allele variable regions to overcome differences in technique (such as primer selection, PCR conditions, and sequencing platform) and successfully combine data from multiple studies into one cohesive dataset. The resultant phylotypes are features that are suitable for the sort of machine learning meta-analysis and personalized medicine that has been successfully deployed for other sorts of ‘omics data to advance mechanistic understanding and treatment outcomes. The phylogenetic placement technique implemented in MaLiAmPi directly facilitates meta-analysis of 16S rRNA gene based studies, overcoming limitations of studies with limited numbers and collecting specimens from participants at a single site, or handful of sites. Further, this is a practical way to relate a specimen from an individual patient to a larger set of observational data that has related the microbiome to outcomes, treatment response, or risk for disease. Thus this approach is a means to activate the microbiome data already collected as a biomarker for precision medicine.

The phylotype-count tables generated by this approach are compatible with approaches like percentile normalization, allowing further integration of microbiome data sets generated by different studies. We have explored integration of 16S rRNA gene data with shotgun metagenomic data, further bolstering the studies that can be integrated.

As we have noted, the technique cannot overcome some fundamental challenges. If the primers selected for the study fail to amplify a critical member of the community, this technique itself cannot infer the presence of those organisms. The lower-read depth of other pyrosequencing based studies result in a limit of detection difference that also cannot be overcome. This limit of detection challenge is shared by approaches like low-read-depth WGS. Further, this approach cannot address technical variance introduced by differences in collection and DNA extraction protocols, with the latter a particularly potent issue when comparing across studies.

This approach also adds a hyperparameter that must be selected: a phylogenetic distance at which to cluster ASVs. For now, we have had good results with phylogenetic distances between 0.1 and 1, but this is a parameter that must be optimized for a given set of studies to be integrated. This can be somewhat mitigated by using phylogenetic distance between communities, (KR-distance^20^) as the feature of interest, eliminating this specific phylotype-clustering hyperparameter.

We believe that phylogenetic normalization of 16S rRNA gene variable region amplicon sequence variants is a promising approach for harmonizing microbiome data from different studies, that significantly outperforms existing techniques such as closed-OTU generation. The outputs are suitable for both meta-analysis and precision medicine. This approach is fully implemented as a reproducible and portable Nextflow-based workflow that we hope will facilitate future microbiome studies.

## Methods

### Phylogenetic placement of 16S rRNA gene ASVs via MaLiAmPi

MaLiAmPi ^30^ (Maximum Likelihood Amplicon Pipeline) is a *Nextflow*-based workflow that implements the approach described in this article. The workflow is 100% containerized and portable, and can be run locally (via Docker), on public clouds (such as Amazon Web Services Batch), or academic high performance computing clusters (e.g. SLURM or Sun Grid Engine-based) via Singularity containers. There are four broad steps MaLiAmPi implements: (1) generation of ASVs; (2) selection of a repository of full-length 16S rRNA alleles; (3) generation of a reference package including a phylogenetic tree of full-length 16S rRNA alleles from the repository that match the ASVs ; and (4) placement of the ASVs onto the reference package phylogenetic tree.

### 1. Generation of amplicon sequence variants (ASVs) from FASTQ files

As noted in the Main section, the overall approach is relatively agnostic to the method used to generate ASVs. MaLiAmPi uses DADA2 by default, based in part on prior benchmarking studies ^31^. For Illumina reads, if index reads are available demultimplexing is confirmed with `Barcodecop` (version 0.5). Reads are then filtered, trimmed and have residual primer and linker sequences removed with `TrimGalore` (version 0.6.6--0). Amplicon sequence variants are then generated using DADA2 (version 1.18.0). Reads are grouped into Batches, ideally representing a group of specimens processed into a library together, and typically of a size of 100.

Each specimen’s reads (or read pairs) are then filtered and trimmed (in parallel) with DADA2’s `filterAndTrim` with the following parameters for Illumina reads:

|  |  |
| --- | --- |
| `maxN` | `0` |
| `maxEE` | `Inf` |
| `truncQ` | `2` |
| `trimLeft` | `0` |
| truncLen | (0, 0) |

And with the following parameters for 454/Pyrosequencing reads:

|  |  |
| --- | --- |
| maxN | 0 |
| maxEE | Inf |
| truncQ | 2 |
| trimLeft | 0 |
| truncLen | 250 |
| maxLen | Inf |

Filtered and trimmed reads are then dereplicated with the DADA2’s `derepFastq` command. The filtered and trimmed reads are grouped into batches, and then the `learnErrors` command is used to generate an error model for each batch’s forward (and when available) reverse reads with the following parameters for Illumina data:

|  |  |
| --- | --- |
| MAX_CONSIST | 10 |
| Randomize | TRUE |
| nbases | 1e8 |

And these parameters for 454/Pyrosequencing data:

|  |  |
| --- | --- |
| MAX_CONSIST | 10 |
| Randomize | TRUE |
| nbases | 1e8 |
| HOMOPOLYMER_GAP_PENALTY | -1 |
| BAND_SIZE | 32 |

By batch, the batch’s error model is applied to the dereplicated reads using the `dada` command with the `pool=“pseudo”` option for all data, additionally `HOMOPOLYMER_GAP_PENALTY=-1, BAND_SIZE=32` for 454/pyrosequencing data.

On a per-specimen basis, paired-end reads are merged with the `mergePairs` command with the following parameters:

|  |  |
| --- | --- |
| trimOverhang | TRUE |
| maxMismatch | 0 |
| minOverlap | 12 |

The `minOverlap` parameter occasionally needs to be relaxed down to a lower number depending on the PCR primer design and specific Illumina chemistry used, specifically when most or all read pairs fail to merge. For very-low quality read data (e.g. when read pairs fail to merge even with a min overlap of 4), we will only use the forward read data (as we believe those reads cannot be accurately paired).

Finally the merged read pairs or dada models for unpaired reads are converted to sequence tables with the `makeSequenceTable` command. From these sequence tables are the ASV sequences and specimen-ASV counts extracted into FASTA and CSV formats respectively for subsequent analysis.

### 2. Repository sequence selection

We started with the deduplicated -> 1200bp -> filtered -> named subset of 16S rRNA alleles from NCBI via the YA16SDB pipeline as our repository of sequences. As noted in the Main section, other repositories of 16S rRNA alleles can also be employed (e.g. SILVA, RDP, greengenes, etc). This entire set of YA16SDB reads are available for download (as below in the Data Availability section) on Zenodo (doi: 10.5281/zenodo.6876634).

A subset of repository candidate full-length 16S rRNA alleles are identified by searching the repository sequences for matches with at least 80% identity to at least one ASV sequence using `vsearch` (version 2.17.0) in `usearch_global` mode, and `max_accepts=10`. To ensure the resultant tree will not result in overfitting or over diffusion of ASV placement later, full-length 16S rRNA alleles are recruited from the repository with the objective of having roughly the same number of recruited reference sequences per each amplicon sequence variant. Specifically, we establish the best possible percent identity between each ASV and the repository alleles, and discard any alleles that are below this best possible percent identity (e.g. retain the bounded-best-hits). We then determine how many ASVs each reference is a best hit for and discard those that are not a best hit for at least two ASVs. Finally we backfill references for ASVs that no longer have a reference sequence as good as their best it, focusing on the longest alleles with no ambiguous bases and with a precise taxonomic annotation. Even for very broad sets of ASVs, this typically results in less than 30,000 reference alleles.

### 3. Reference package recreation

These filtered reference alleles are now aligned with `cmalign` from the `Infernal` package using the `SSU_rRNA_bacteria` covariance matrix from the `rfam` database and a `mxsize 4096`. The recruited full-length 16S rRNA alleles alignment is then assembled into a phylogeny. The generation of the phylogenetic tree is the most computationally intensive step in the entire approach. The current implementation default to `RAxML (version 8.2.4)`, but also allows `RAxML-ng` (1.0.3) to be used if desired for a deeper exploration of possible starting random trees.

For `RAxML`, the following settings are used:

|  |  |
| --- | --- |
| -m | GTRGAMMA |
| -p | 12345 |

And for `RAxML-ng`:

|  |  |
| --- | --- |
| model | GTR+G |
| seed | 12345 |
| tree | pars{1},rand{1} |
| bs-cutoff | 0.3 |

This *de novo* phylogenetic tree is combined with the metadata for each allele within the tree (e.g. species-level taxonomy, source accession, etc) into a standardized reference package format using the `taxtastic` package.

### 4. Placement of ASVs onto a reference package phylogenetic tree

ASVs are next placed onto this reference tree. First the ASV sequences are aligned, using `cmalign` from the `Infernal` package, and the same covariance matrix as was used to make the alignment of reference sequences (retained in the reference package). The ASV alignment is combined with reference alignment (contained within the reference package) using `esl-alimerge` utility from `easel`.

This combined alignment is then used to phylogenetically place the ASVs onto the reference package tree using either `pplacer` (the current default) or `epa-ng`. Both have comparable performance and outputs. For `pplacer`, the following parameters are used:

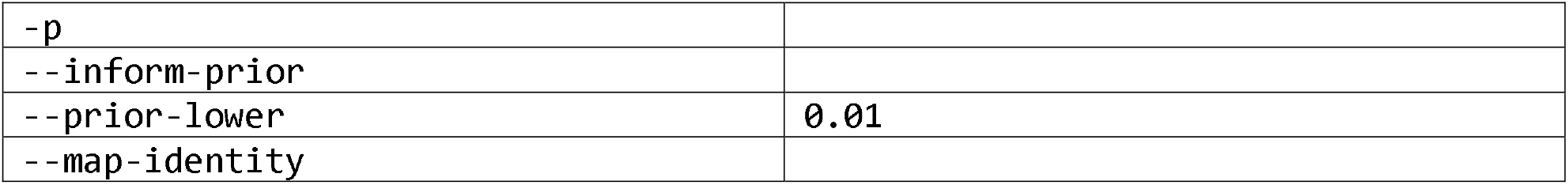

|  |  |
| --- | --- |
| `-p` |  |
| `--inform-prior` |  |
| `--prior-lower` | `0.01` |
| `--map-identity` |  |

For epa-ng:

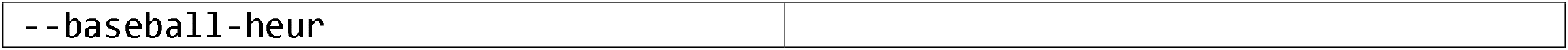

The output of the placement step is in JPLACE format, `dedup.jplace`. For each ASV, the likelihood, distal-length, and pendant-length is reported for each edge in the tree (omitting edges for which there is no meaningful likelihood). These likelihood-weighted trees are the basis for subsequent analysis. Combined with ASV-counts-per-specimen, the weighted tree can be used to estimate pairwise phylogenetic distance (KRD-distance, akin to weighted UniFrac) between specimens, the alpha diversity of a specimen, and to group ASVs into phylotypes. Phylotypes are groups of ASVs clustered at a specific phylogenetic distance, and are created using a Python package (https://github.com/jgolob/phylogroups) installable via pypi (https://pypi.org/project/phylotypes/). A distance of 1 roughly corresponds to a species of bacteria, but with significant variation depending on the degree of taxonomic - phylogenetic concordance.

### In silico human gut microbiota for validation

As in our prior work^18^, we used 100 microbial communities similar in structure and composition to those found in the healthy human gut microbiome, but generated *in silico* and thus with a known allele of origin for each and all amplicons generated. These communities are available via Zenodo (10.5281/zenodo.1120359). For each community, we have selected specific full-length unambiguous 16S rRNA gene alleles to represent each microbe within the community. From these alleles we can generate amplicons targeting specific hypervariable regions via *in silico* PCR.

We selected primers targeting the most common variable domains and sequencing platforms represented in the large volume of legacy 16S rRNA gene data available in public repositories. Specifically, the V4 region (or V3-V6), V1-V2, and V5-V9 domains (**Figure 1**) and the sequencing platforms Illumina MiSeq or Roche 454 (a legacy technology for which SRA contains 139,965 records with the label ‘16S’). For MiSeq we set a goal of 50,000 simulated amplicons per community and for 454 we targeted 5,000 amplicons per community, reflecting the typical read-depths from the respective platforms (**Table 2**). As depicted in **Figure 1**, there is effectively no overlap between the amplicons targeting distinct regions (i.e., no overlap in sequence between the primers targeting V1-V2 and V5, nor with V6-V9).

**Table 2:** Primers for amplification of 16S rRNA gene variable regions and subsequent sequencing that were evaluated in this study. For MiSeq primers, we simulated 50,000 amplicons per community; for 454, we simulated 5,000 amplicons per community.

| Primer Set | Variable Region | Intended Platform | Simulated amplicons per community |
| --- | --- | --- | --- |
| 27fmod / 338r | V1 - V2 | Illumina MiSeq | 50,000 |
| U515f / 806r | V4 | Illumina MiSeq | 50,000 |
| 27f / 357r | V1 - V2 | 454 | 5,000 |
| 357f / 926r | V3 - V5 | 454 | 5,000 |
| 968f / 149r | V6 - V9 | 454 | 5,000 |

### Dereplication of ASVs

ASVs with the exact same sequence (length and each base pair) were combined together and assigned an ID.

### Generation of closed OTUs

Here we used the QIIME1 package, and the GreenGenes 97% OTU repository. We generated a docker container containing QIIME 1 version 1.9.1A, and ran the following commands to generate blast-picked closed OTUs with a similarity of at least 80%:

pick_otus.py -i <raw_fastq> -o blast_picked_otus/ -m blast -r 97_otus.fasta -s 0.8

Where the 97_otus.fasta were the 97_otus from the GreenGenes repository, as recommended by the QIIME1 documentation.

### Calculation of Bray-Curtis distance

Count tables were first assembled with one row per specimen and one column per feature (dereplicated ASV, closed-OTU, or phylotype) and each cell the number of reads assinged to that feature and specimen. These raw-count tables were then normalized to a read depth of 10,000 reads per specimen. The normalized count tables were then used to calculate pairwise Bray-Curtis distance using the *scipy* (verison 1.6.3) pairwise distance calculator.

### UMAP ordination

The *Python* umap-learn package (version 0.5.1) was used with the following hyperparameters:

min_distance = 0

n_components = 2

n_neighbors = 45

Random state was fixed at 42. The pre-computed Bray-Curtis distance (as above) was used.

### Correlation between ‘real’ and ‘estimated’ pairwise Bray-Curtis distance

For the real pairwise distance between communities, the distances between communities determined using the *same* primers were used. For estimated, the distance between the same community, but with each community amplified with a *different* primer was used.

### Clustering via HDBSCAN

Pairwise Bray-Curtis distance was calculated from normalized phylotype-counts as described above. This pairwise distance matrix was used for ordination with umap, with the following hyperparamters:

min_distance = 0

n_neighbors = 100

n_components = 2

The python hdbscan package was used, using the ordinated points per specimen as the input matrix and the following hyperparameters:

min_cluster_size = 25

min_samples = 2

## Data Availability

The in silico data sets used are available via Zenodo, at 10.5281/zenodo.1120360

The set of full-length reference 16s rRNA alleles can be found on Zenodo at 10.5281/zenodo.6876633.

Real-world human microbiome data is available on the NCBI Sequence Read Archive (SRA) under the BioProjects in **Table 1**.

## Code Availability

MaLiAmPi is available via a git repository (https://github.com/jgolob/maliampi).

ARF is a workflow used to create the repository of full-length 16s rRNA alleles. It is available as a git repository (https://github.com/jgolob/arf).

## Acknowledgements

The authors would like to thank Teresa O’Meara and Thomas Schmidt for their expert edits of the manuscript and text.

