## Supplemental Figures and legends for "Robust Harmonization of Microbiome Studies by Phylogenetic Scaffolding with MaLiAmPi"

**FIGURE S1**

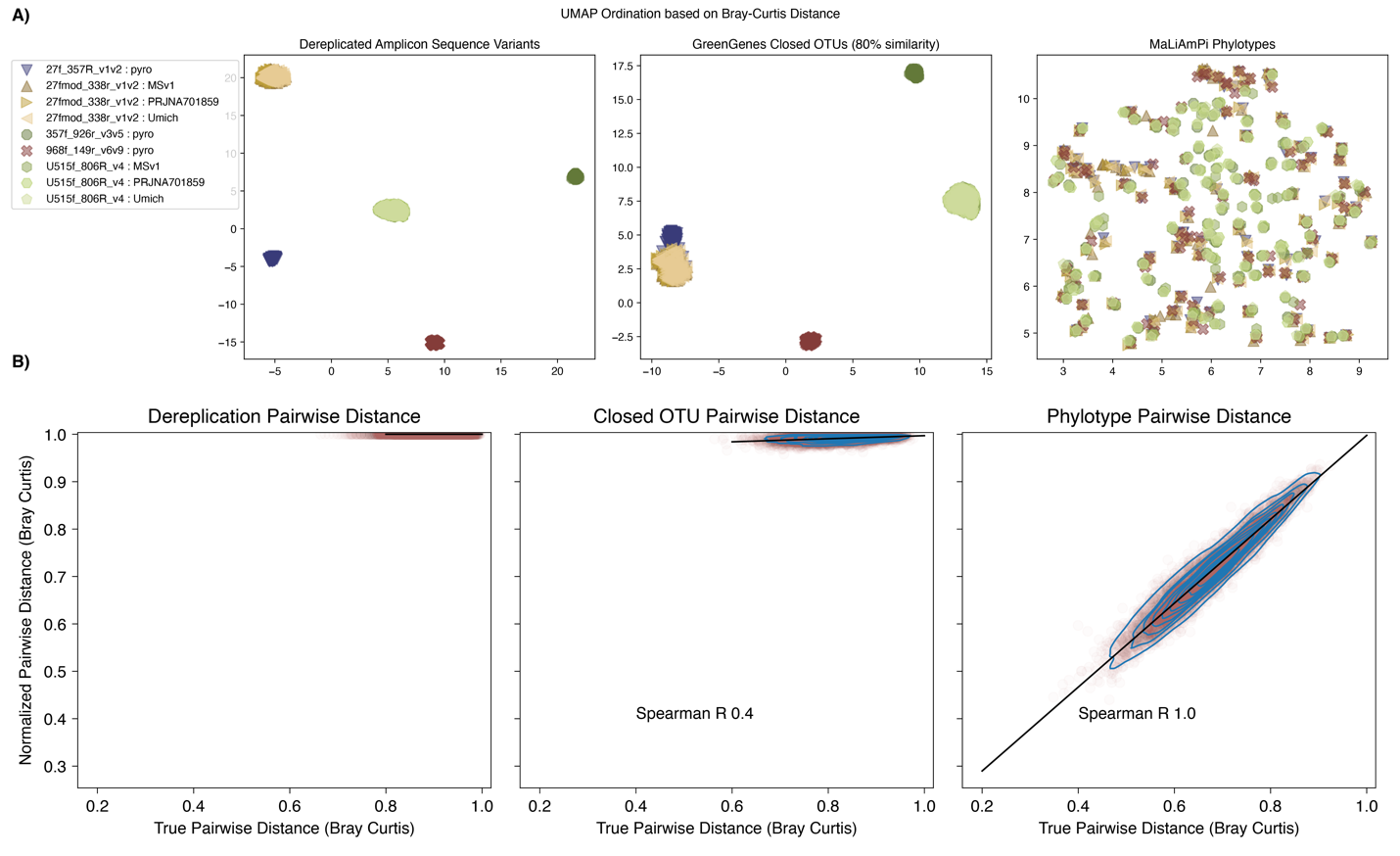

**Figure S1: Successful integration of 16S rRNA amplicons targeting different variable regions and with sequencer-induced error after phylogenetic placement.** UMAP Ordination Plots based on Bray-Curtis distance between specimens, as determined from count-tables based on DADA2 ASVs (left), Closed-reference OTUs (center), or phylotypes derived from phylogenetic placement (right). A) are all 100 communities, with different markers for each primer set. As expected, DADA2 ASVs were able to correctly combine reads generated with the same primers but different sequencer error rates but not reads generated with different primers amplifying distinct regions of the 16S rRNA gene. Closed reference OTUs were able to overcome differences in sequencer, but not variable region. In contrast, phylotypes were able to integrate data generated from the same underlying 100 microbial communities, but observed by amplifying distinct variable regions and using different sequencers with different error rates. B) shows the correlation between ‘true’ pairwise distance between microbial communities (as estimated when using the same primer and error models) as compared to estimated distance after normalization of the same community pairs, but now as sequenced with different primers.

**FIGURE S2**

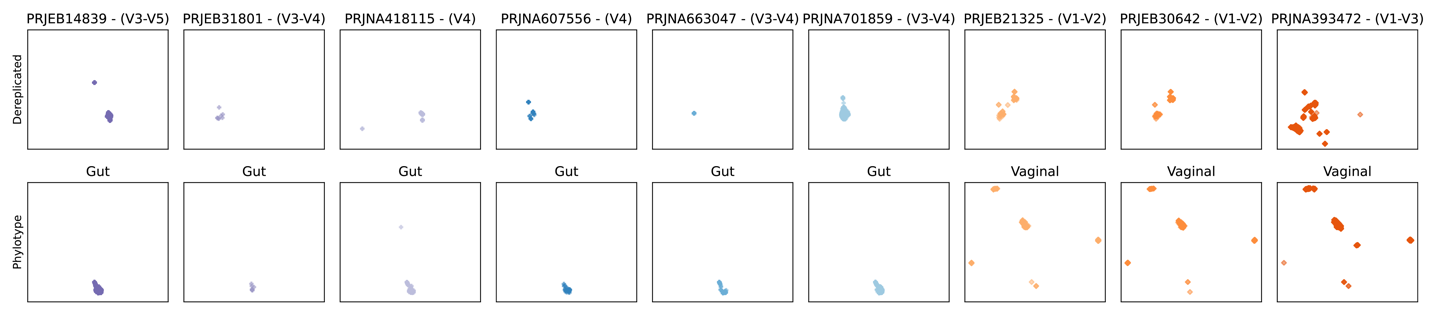

**Figure S2: UMAP plots based on Bray-Curtis distance from dereplicated ASV counts (top) or phylotype counts (bottom) stratified by project**.
